# Amazonian soil metagenomes indicate different physiological strategies of microbial communities in response to land use change

**DOI:** 10.1101/2020.09.15.299230

**Authors:** Md Abdul Wadud Khan, Brendan J. M. Bohannan, Kyle M. Meyer, Ann M. Klein, Klaus Nüsslein, James P. Grover, Jorge L. Mazza Rodrigues

## Abstract

Despite the global importance in ecological processes, the Amazon rainforest has been subjected to high rates of deforestation, mostly for pasturelands, over the last few decades. In this study, we used a combination of deep shotgun metagenomics and a machine learning approach to compare physiological strategies of microbial communities between contrasting forest and pasture soils. We showed that microbial communities (bacteria, archaea and viruses), and the composition of protein-coding genes are distinct in each ecosystem. The diversities of these metagenomic datasets are strongly correlated, indicating that the protein-coding genes found in any given sample of these soil types are predictable from their taxonomic lineages. Shifts in metagenome profiles reflected potential physiological differences caused by forest-to-pasture conversion with alterations in gene abundances related to carbohydrate and energy metabolisms. These variations in these gene contents are associated with several soil factors including C/N, temperature and H^+^+Al^3+^ (exchangeable acidity). These data underscore that microbial community taxa and protein-coding genes co-vary. Differences in gene abundances for carbohydrate utilization, energy, amino acid, and xenobiotic metabolisms indicate alterations of physiological strategy with forest-to-pasture conversion, with potential consequences to C and N cycles. Our analysis also indicated that soil virome was altered and shifts in the viral community provide insights into increased health risks to human and animal populations.

## IMPORTANCE

In the Amazon, forest-to-pasture conversion has a strong negative impact on biological composition that results in alteration of important ecosystem processes. As the most comprehensive survey known to date, this study with shotgun metagenomics approach demonstrates predictive understanding of how an anthropogenic activity shapes microbial taxonomy and functions across ecosystems, and the link between them. We then link the microbiomes with each soil type to explore the adaptation mechanisms and ecological traits of microbial communities in their ecosystems. Among these, carbon metabolism and energy harvest strategies mostly relate to specific environmental challenges of microbial communities, providing insights to understand the mechanisms of ecologically relevant metabolic processes. Despite their critical roles in biogeochemical processes, viruses in terrestrial ecosystem remains understudied. Thus, the inclusion of viral community in this study would shed new lights on their roles in microbial functional traits, and provide novel hypotheses in microbial ecology of terrestrial ecosystem.

## INTRODUCTION

The Amazon is the largest continuous rainforest on Earth and provides essential ecosystem services at regional and global scales. It harbors the largest collection of plants and animal species in the world (1) and balances the flux of atmospheric gases (2). Despite these known benefits, forest clearing has proceeded at an alarming rate over the last few decades. The process of deforestation and land use change concurrently alters the biological and biochemical compositions of the soil (3–6), which could have important consequences to the associated ecosystem processes.

Land use change is predicted to be the most important factor in altering biodiversity in tropical areas for the twenty first century (7). Ecologists have long studied the contribution of plant and animal communities to ecosystem functions in tropical forests (8–10) and the impact of deforestation on these communities (3, 4). In contrast, microbial ecologists have just started to explore similar sets of questions in the Amazon, showing that ecosystem conversion alters bacterial (5, 11–13), archaeal (14–16) and fungal (17) community composition in various Amazonian land use types. While some studies showed that plant diversity increases overall microbial diversity (18, 19) and activity in soil (20), others demonstrated the consequences of deforestation were restricted to specific functional groups of microbes or specific protein-coding genes such as those associated with C and N cycles (14, 21–26). The above studies, however, largely relied on either 16S rRNA or a single functional gene as markers of microbial communities. Although, metagenomics approach has been used in few recent studies to examine the effect at two and four months (27) and of forest-to-agriculture conversion (28) following deforestation. However, these metagenomics studies did not either evaluate the long-term impact of deforestation or functional capacities of the most common land use type following deforestation (i.e., pasture) on ecological processes carried out by soil microorganisms.

In addition, viruses remain neglected in Amazon soils even though they are the most abundant biological entity, with an estimated 10^31^ on the planet (29). Viruses not only serve as the largest reservoir of genetic material (30–32), but also play an important role in mediating lateral gene transfer (33) and controlling microbial populations (34). Despite this, very few empirical studies have been conducted on soil viromes (Graham et al. 2019; Williamson et al. 2017), and none in the context of deforestation in the Amazon rainforest.

To examine the effect of forest-to-pasture conversion, the most common land use type following deforestation in the Amazon rainforest, we collected soils samples from pristine forest and 38 years old pastureland, and performed deep sequencing with whole genome shotgun approach, aiming to: (*i*) estimate the composition of microorganisms and protein-coding genes, and their relationships in primary forest and established pasture; (*ii*) identify differential abundances of key protein-coding genes involved in ecologically relevant biochemical processes; and (*iii*) examine the Amazon soil virome and gain a comprehensive understanding of the alterations happening with deforestation.

## RESULTS

### Metagenome profiles

We obtained a mean of about 636 million (±12%) reads per sample, totaling 6.4 billion paired-end reads with an average length of 171 bp. Following quality filtering of metagenomes through MG-RAST (version 3.3.6), the sequence reads were mapped to the KEGG (Vol. 36) and RefSeq (Vol. 37) databases, where each read was annotated according to the KEGG Orthology groups (KOs) and species (bacteria, archaea and viruses) associated with the identified alignments. While the vast majority of the sequence reads (83.7%) passed the quality control by MG-RAST, only 13.4% had protein-coding (KOs) and taxonomic annotations. From KO sequencing reads ranging 60,373,709-76,838,373; taxonomic reads of bacteria ranging 783,394-1,143,853, archaea ranging 20,661-51,138; and viral reads ranging 18,307-32,903, we obtained a total number of unique 5,607 KO genes; 23,949 bacterial, 706 archaeal, and 1,774 viral taxa at species level. Before downstream analyses, all samples were rarefied to their lowest sequencing depth to bring them to equal values in order to remove sequencing effort bias. The rarefaction plots showed that diversity levels of these metagenomic datasets tend to saturate, suitable for a comprehensive analysis of the microbial communities and their physiological potentials in both land use types (**Fig. SF1**).

### Impact of land use change on bacterial and archaeal communities

We observed substantial variation of major bacterial and archaeal taxa between forest and pasture soils. With over 26% [±0.82%; standard error of mean (S.E.M.)] in forest and 19% (±0.22%; S.E.M.) in pasture (*P* < 0.001), *Alphaproteobacteria* was the dominant bacterial class in both ecosystems and showed the largest variation in response to forest-to-pasture conversion (**Fig. SF2A**). Other major bacterial groups that differed significantly between forest and pasture (*P* < 0.001) include: *Actinobacteria* and *Firmicutes*, which increased from 13.1% (±0.21%; S.E.M.) to 16.65% (±0.41%; S.E.M.) and 11.39% (±0.52%; S.E.M.) to 14.4% (±0.17%; S.E.M.), respectively. *Euryarchaeota* and *Thaumarchaeota*, were the two most differentially abundant archaeal phyla (*P* < 0.001), where the former increased from 39.3% (±0.59%; S.E.M.) to 55.48% (±0.52%; S.E.M.), and latter decreased from 27.41% (±0.47%; S.E.M.) to 7.73% (±0.72%; S.E.M.) following deforestation (**Fig. SF2B**). We also identified differentially abundant bacterial (**Table ST1A**) and archaeal (**Table ST1B**) species in each ecosystem. Using criteria set by DESeq2 and Random Forest methods in this study, we identified 131 bacterial species that were enriched in forest soils, and 189 species that were enriched in pasture soils. *Bradyrhizobium* was the most dominant bacterial genus in the forest soils, whereas *Bacillus* and *Arthrobacter* dominated pasture soils. With the same criteria, 101 archaeal species were estimated to be differentially abundant in forest, where an unclassified crenarchaeotal genus was the most dominant. On the other hand, pasture soils had 51 species that were differentially abundant with an unclassified euryarchaeal genus being the most dominant.

### Diversity patterns of taxonomic lineages and protein-coding genes

Forest-to-pasture conversion increased alpha [*P* < 0.01, (**Fig. 1A**)] but decreased beta [*P* < 0.05 (**Fig. 1B**)] diversities of bacterial species. While archaeal diversity did not follow a similar alpha diversity pattern (**Fig. 1C**), beta diversity was statistically different between forest and pasture [*P* < 0.05, (**Fig. 1D**)]. On the other hand, diversities of KO genes resembled those of bacteria [alpha: *P* < 0.05, (**Fig. 1E**), and beta: *P* < 0.01, (**Fig. 1F**)]. We then asked whether the above bacteria and archaea diversity results co-varied with protein-coding genes diversity values. In this analysis, we used the principal coordinate loadings of the first axis (PCo1), which explained the highest variation in the metagenome compositions. Our data showed strong compositional similarities of KO gene profiles with bacterial [*R*^2^ = 0.95, *P* < 0.05 (**Fig. 2A**)] and archaeal [*R*^2^ = 0.93, *P* < 0.05 (**Fig. 2B**)] species profiles.

**FIG. 1.**
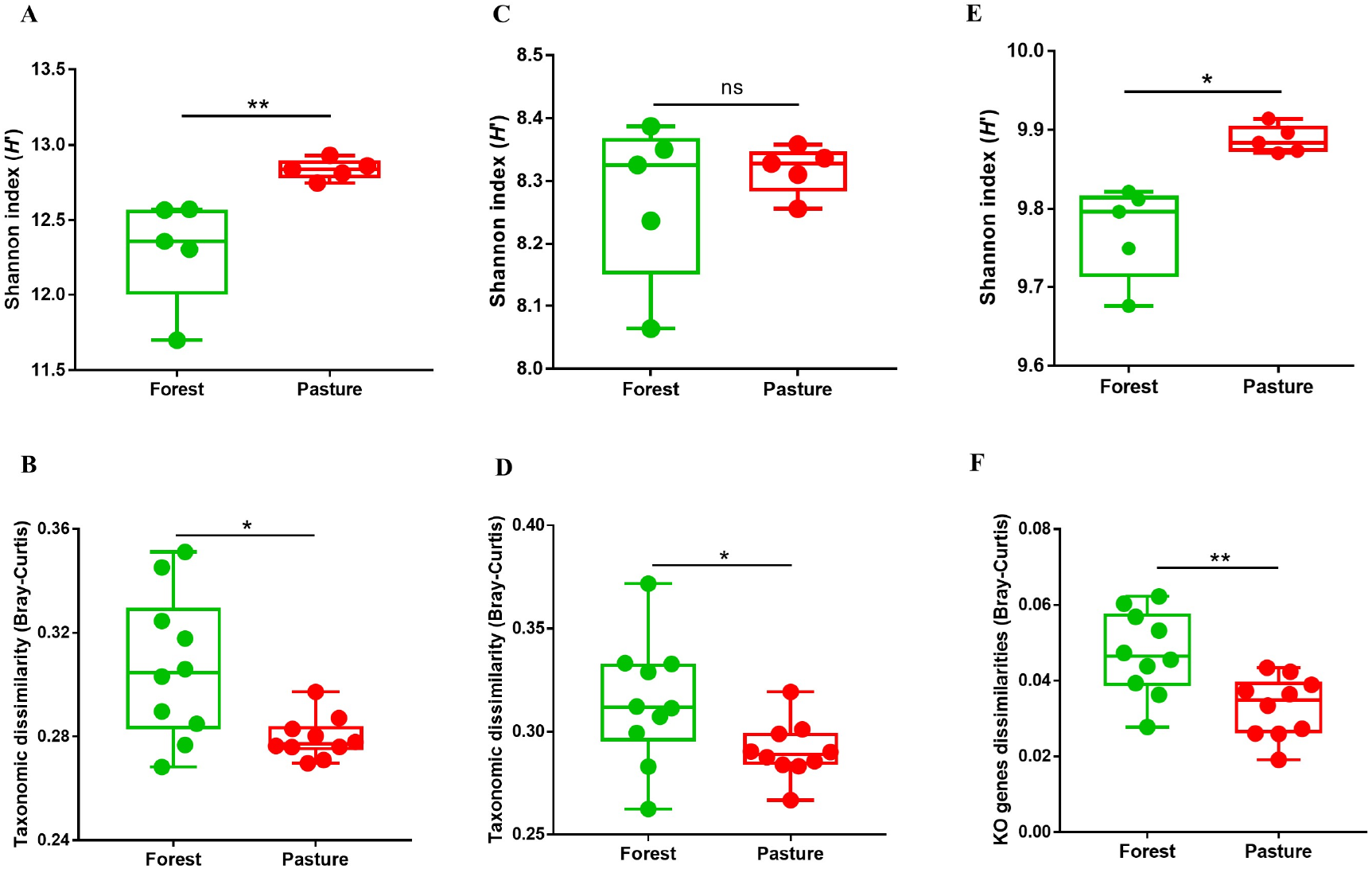
Response of forest-to-pasture conversion to bacterial (A-B), archaeal (C-D) and protein-coding gene (E-F) alpha and beta diversities in soil metagenomes. Shannon index and Bray-Curtis similarity were used for estimating alpha (n = 5), and beta diversities (n = 10 for pairwise similarity), respectively, where mean values of diversity were depicted. Error bars represent standard error of the mean (S.E.M). Statistical significance was calculated using Mann-Whitney test with 1,000 permutations. Symbols *, and ** indicate significance values of *P* <0.05 and *P* < 0.01, respectively.

**FIG. 2.**
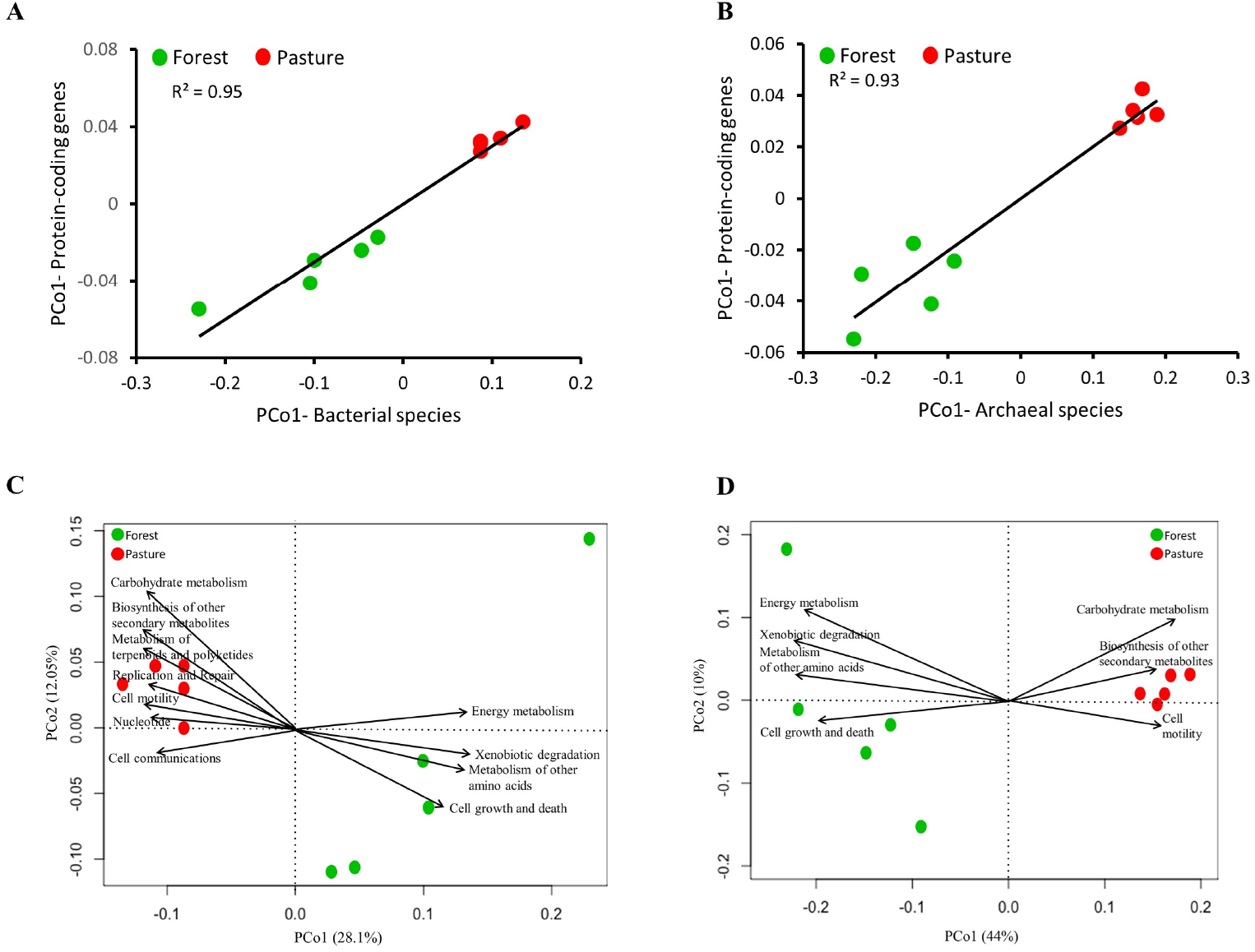
Relationships of protein-coding gene compositions with bacterial and archaeal species compositions across forest and pasture metagenomes. The relationships of compositional similarities are visualized using the principal coordinate loadings for the first axes (PCo1). Protein-coding genes demonstrated significantly positive correlations with bacterial (A) and archaeal (B) community compositions. Vector fitter principal coordinate analysis showed that bacterial (C) and archaeal (D) communities have distinct clustering patterns by soil types, where vectors represent the association of the functional characteristics at KEGG level 2. The vectors are shown with arrows pointing to the direction of an increase in the gradient for the corresponding variable, and with length proportional to the correlation between ordination and variable. The circles (forest, green; pasture, red) represent the relative positions of each soil taxonomic community. The significances (*P*-values) of the vectors were calculated based on 999 random permutations of the data. Here, only functional characteristics that were estimated to be significantly (*P* < 0.01) associated with PCo1 are shown. Bray-Curtis metric was used for estimating distances between different samples.

### Impact of land use change on major functional characteristics

The ordination plots based on principal coordinate analysis (PCoA) showed that the taxonomic composition in the forest soil clustered apart from that in the pasture [ANOSIM: *R* = 0.74, *P* = 0.007 and *R* = 0.99, *P* = 0.006, respectively (**Fig. 2C and 2D**)]. Next, we fit KEGG level 2 functions as vectors onto the ordination and several functional attributes were significantly associated with the bacterial PCo1 (28.1%) across different soil samples (**Fig. 2C**). Positive correlations of PCo1 with energy metabolism (r^2^ = 0.92, *P* < 0.001), cell growth and death (r^2^ = 0.98, *P* < 0.001), xenobiotics biodegradation and metabolism (r^2^ = 0.96, *P* < 0.001), and metabolism of other amino acids (r^2^ = 0.96, *P* < 0.01), as indicated by the direction of the arrows, demonstrated that increasing gradients of these functions were strongly associated with forest samples, whereas pasture samples were located at opposite ends of these gradients. Similarly, pasture sites had increasing gradients of carbohydrate metabolism (r^2^ = 0.86, *P* < 0.01), cell motility (r^2^ = 0.89, *P* < 0.001), metabolism of terpenoids and polyketides (r^2^ = 0.58, *P* < 0.05), nucleotide (r^2^ = 0.67, *P* < 0.05), cell communications (r^2^ = 0.71, *P* < 0.05) and biosynthesis of other secondary metabolites (r^2^ = 0.91, *P* < 0.01), whereas forest sites were at decreasing gradient of them. Most of these functional traits were also estimated to be associated with the archaeal species (**Fig. 2D**), where strength and direction of associations were alike. Similar trends were also observed when performing a comparison for relative abundances of functional categories at KEGG level 2 between land use types (**Fig. SF3**).

### Differential abundance of protein-coding genes and metabolic pathways

Next, we focused on determining what metabolic pathways (KEGG level 3) and associated protein-coding genes (KEGG level 4) were responsible for the disparities observed in major functional categories (KEGG level 2) in relation to land use change. A comparative analysis identified 935 KO genes showing large differences between forest and pasture metagenomes, of which 541 belong to KEGG Enzyme Commission (EC) categories. A complete list of KO genes with log_2_FC values and the *P*-values of Wald tests is provided in **Table ST2**. Our analysis showed that the major differences of forest and pasture metagenomes were associated with genes involved in carbohydrate, energy, and amino acid metabolisms and xenobiotic degradation, which are functionally connected to other pathways (**Fig. 3**).

**FIG. 3.**
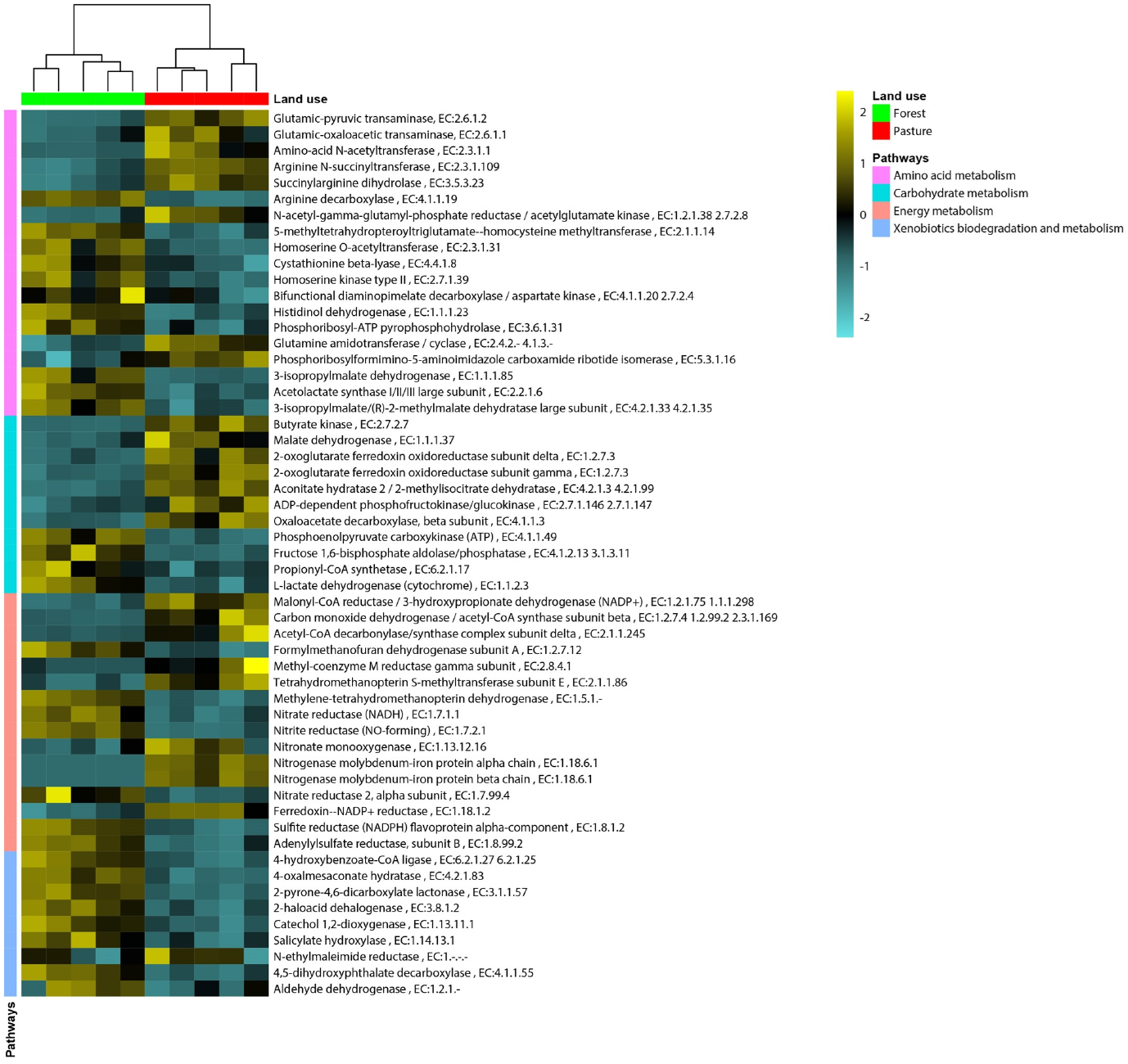
Heat map comparison of *Z*-scores calculated from protein-coding gene abundances between forest and pasture metagenomes. Only subset of genes encoding KEGG ECs involved in nutrient cycling is shown and a complete list of all discriminatory protein-coding genes is shown in **Table ST2**. Only genes that passed both selection criteria for DESeq2 (BH-adjusted *P* < 0.05) and Random Forest (importance score > 0.0001) methods were considered.

#### Carbohydrate metabolism

Pasture soil samples were estimated to contain elevated numbers of KEGG EC genes associated with the metabolism of starch, sucrose, and galactose (**Table ST2**). In addition, KO genes involved in the transport of mono- and di-saccharides were far more abundant in pasture, of which log_2_FC values ranged from −0.39 to −3.01. Unlike pasture metagenomes, which had elevated counts of glycolysis-specific genes [e.g., ADP-dependent phosphofructokinase/glucokinase (EC:2.7.1.146, EC:2.7.1.147)], forest metagenomes had higher counts of genes specific for gluconeogenesis [e.g., fructose 1,6-bisphosphate phosphatase (EC:3.1.3.11); **Fig. 4**). In addition, genes associated with the pentose phosphate pathway (PPP) and the conversion of oxaloacetate to phosphoenolpyruvate [PEP; phosphoenolpyruvate carboxykinase (EC4.1.1.49)] had higher relative abundances in the forest metagenome samples. Furthermore, we observed considerable variation between forest and pasture metagenomes in frequencies of genes that are involved in succeeding metabolism of sugars. For example, genes associated with the tricarboxylic acid (TCA) cycle were estimated to be higher in pasture soils (**Fig. 4**). In addition, pasture soils had higher abundances of genes affiliated with fermentation of acetate (e.g., acetyl-CoA hydrolase (EC:3.1.2.1), and butanoate [e.g., butyrate kinase (EC:2.7.2.7)], whereas genes associated with fermentation of propanoate [propionyl-CoA synthetase (EC:6.2.1.17)], and lactate [L-lactate dehydrogenase, cytochrome (EC:1.1.2.3)] were elevated in forest soils.

**FIG. 4.**
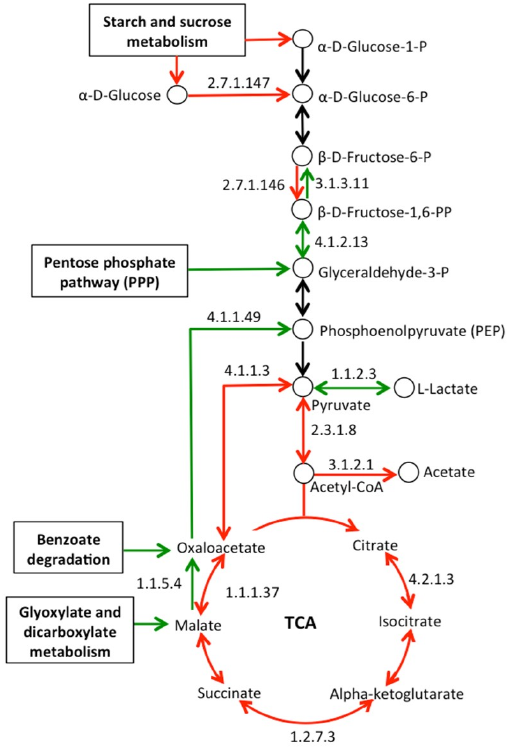
Metabolic map for glycolysis/gluconeogenesis and the tricarboxylic acid (TCA) cycle based on differentially abundant genes observed in forest and pasture metagenomes. Only genes that passed both selection criteria for DESeq2 (BH-adjusted *P* < 0.05) and Random Forest (importance score > 0.0001) methods are represented. Arrows indicate the enzyme-mediated steps of the pathway, with KEGG EC numeric classification representing the reaction being catalyzed. A green arrow means that the abundance of a gene encoding KEGG EC enzyme was higher in the forest, while red means higher in the pasture. Black arrows indicate genes with similar abundances between forest and pasture metagenomes.

#### Energy metabolism

The most prominent differences observed were related to the genes associated with nitrogen and methane metabolisms (**Fig. 5A**). Forest metagenomes had an overrepresentation of EC genes involved in nitrification [nitronate monooxygenase (EC:1.13.12.16); nitrate reductase 2 alpha subunit (EC1.7.99.4)], denitrification [nitrite reductase, NO-forming (EC:1.7.2.1)] and assimilatory nitrate reduction [nitrate reductase, NADH (EC:1.7.7.1)] (**Fig. 5A**). Conversely, the estimated representation of genes involved in nitrogen fixation was higher in the pasture metagenomes [nitrogenase molybdenum-iron protein alpha, and beta chains (EC:1.18.6.1) with log_2_FC values of −5.76 and −5.92, respectively]. In methane metabolism, several genes involved in methanogenesis were enriched in pasture metagenomes [e.g., carbon monoxide dehydrogenase (EC:1.2.7.4); acetyl-CoA decarbonylase/synthase complex subunit delta (EC:2.1.1.245); tetrahydromethanopterin S-methyltransferase (EC:2.1.1.86); and methyl-coenzyme M reductase gamma subunit (EC:2.8.4.1)]. Other genes involved in methanotrophy were enriched in the forest [methylene-tetrahydromethanopterin dehydrogenase (EC:1.5.1.-); formylmethanofuran dehydrogenase subunit A (EC:1.2.7.12)] (**Fig. 5B and Table ST2**).

**FIG. 5.**
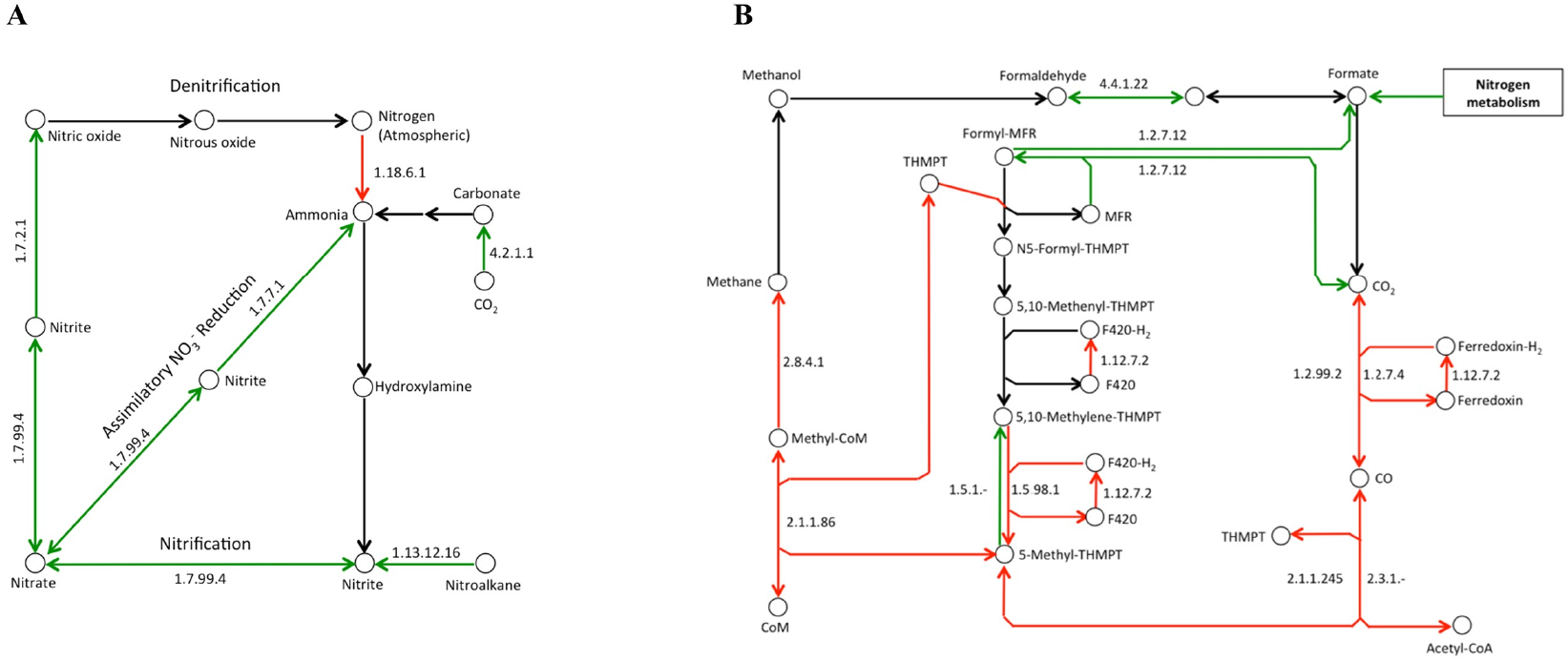
Metabolic maps for the nitrogen (A) and methane (B) metabolisms based on differentially abundant genes observed in forest and pasture metagenomes. Only genes that passed both selection criteria for DESeq2 (BH-adjusted *P* < 0.05) and Random Forest (importance score > 0.0001) methods are represented. Arrows indicate the enzyme-mediated steps of the pathway, with KEGG EC numeric classification representing the reaction being catalyzed. A green arrow means that the abundance of a gene encoding KEGG EC enzyme was higher in the forest, while red means higher in the pasture. Black arrows indicate genes with similar abundances between forest and pasture metagenomes.

#### Amino acid metabolism

The overall abundance of genes involved in amino acid metabolism was not statistically different between forest and pasture. However, we observed overrepresentation of certain genes in forest metagenomes (*P* < 0.05), especially those encoding for enzymes involved in key steps of oxaloacetate-derivative amino acids such as methionine, threonine (**Fig. SF4A**), PPP-derivative amino acids including histidine (**Fig. SF4B**), and pyruvate derivative branched-chain amino acids (valine, leucine and isoleucine) (**Fig. SF4C**). In addition, degradation genes for branched-chain amino acids were also higher in forest metagenomes (**Table ST2**). As a contrast, pasture metagenomes were associated with the metabolism of alpha-ketoglutarate, another TCA cycle intermediate (**Fig. 4**). In addition, there was increased gene abundance related to the biosynthesis of arginine from glutamate, proline from arginine (**Fig. SF4D**).

#### Xenobiotics biodegradation and metabolism

We determined higher gene sequence counts (*P* < 0.05) in forest metagenomes associated with the degradation of diverse aromatic compounds including benzoate, chlorocyclohexane and chlorobenzene, dioxin, nitrotoluene, and polycyclic aromatic hydrocarbon (**Fig. SF5**). These pathways seem to be linked and contribute to the production of the pyruvate and oxaloacetate that might act to support the TCA cycle and replenish its intermediates, a mechanism known as anaplerosis, which are used in the biosynthesis of essential compounds such as amino acids.

#### Others

Pastures had a higher abundance of several genes affiliated with chemotaxis, flagellar assembly, and regulation of the actin cytoskeleton (**Table ST2**). In addition, we estimated several differentially abundant genes between forest and pasture soils that were associated with plant-microbe and microbe-microbe interactions. For example, genes associated with biosynthesis of other secondary metabolites such as of benzoxazinoid, which are plant secondary metabolites, typically from grass species, and a beta-lactam resistance gene observed to be higher in pasture metagenomes. In contrast, a streptomycin synthesis gene encoding for a streptomycin-6-phosphatase (EC:3.1.3.39) was higher in forest metagenomes. Within the two-component system pathway, the abundance of genes for multidrug transport proteins (K07788, K07789) was greater in forest, while bacitracin resistance response regulator proteins (K11629, K11630) were estimated to be higher in pasture metagenomes. Several genes (K13448, K13429, K13462, K13428, K13447 and K13424) that are known to be associated with plant-pathogen interaction pathways were more abundant in pasture metagenomes, while none were detected in forest metagenomes.

### Relationship between the KO gene composition and soil physicochemical properties

To explore potential drivers of community genomic composition, we performed an environmental fitting of 22 soil variables onto an ordination plot via PCoA. KO genes clustered by land use types [(ANOSIM: *R* = 0.98, *P* = 0.007 (**Fig. SF6**)]. Negative correlations of PCo1 with Carbon/Nitrogen (C/N) (r^2^=0.95, *P* < 0.001), temperature (r^2^=0.97, *P* < 0.001), base saturation (V; r^2^=0.78, *P* < 0.01), and Fe (r^2^=0.72, *P* < 0.01) demonstrate that pasture samples have increasing gradients of these factors, whilst forest samples are at opposite ends of these gradients. Similarly, forest samples have increasing gradients of exchangeable acidity (H^+^+Al^3+^), Sulfur (S), and Boron (B), whereas pasture samples are opposite to them.

### Impact of land use change on viral composition and diversity

While evaluating the impact of land use change to genomic content of microbial communities, we observed a large number of reads associated with viral sequences and asked whether land use change impacted the soil virome. We observed that alpha diversity of viral community is higher in pasture compared to forest (**Fig. 6A)**, with significant compositional differences in these two contrasting ecosystems (ANOSIM: *R* = 0.67, *P* = 0.008; **Fig. 6B**). We compared the viral populations at the family level between the two land use types and estimated that *Siphoviridae, Podoviridae*, an unclassified family, and *Myoviridae* were most abundant groups, collectively comprised over 80% and 90% of the sequences found in pasture and forest soils, respectively (**Fig. 6C**). While the relative abundance of sequences for the above families did not differ between land uses, we observed significant differences for several viral families, such as *Bicaudaviridae*, *Microviridae, Caulimoviridae, Tectiviridae* and *Flaviviridae*. We identified 25 viral species that were enriched in forest soils, and 67 species that were enriched in pasture soils (**Table ST3**). In addition, we observed significant compositional associations of viral species with bacteria [(*R*^2^=0.84, *P* < 0.05], archaea [*R*^2^=0.93, *P* < 0.05], and KO genes [*R*^2^=0.86, *P* < 0.05] profiles (**Figs.S7A, S7B, and S7C**).

**FIG. 6.**
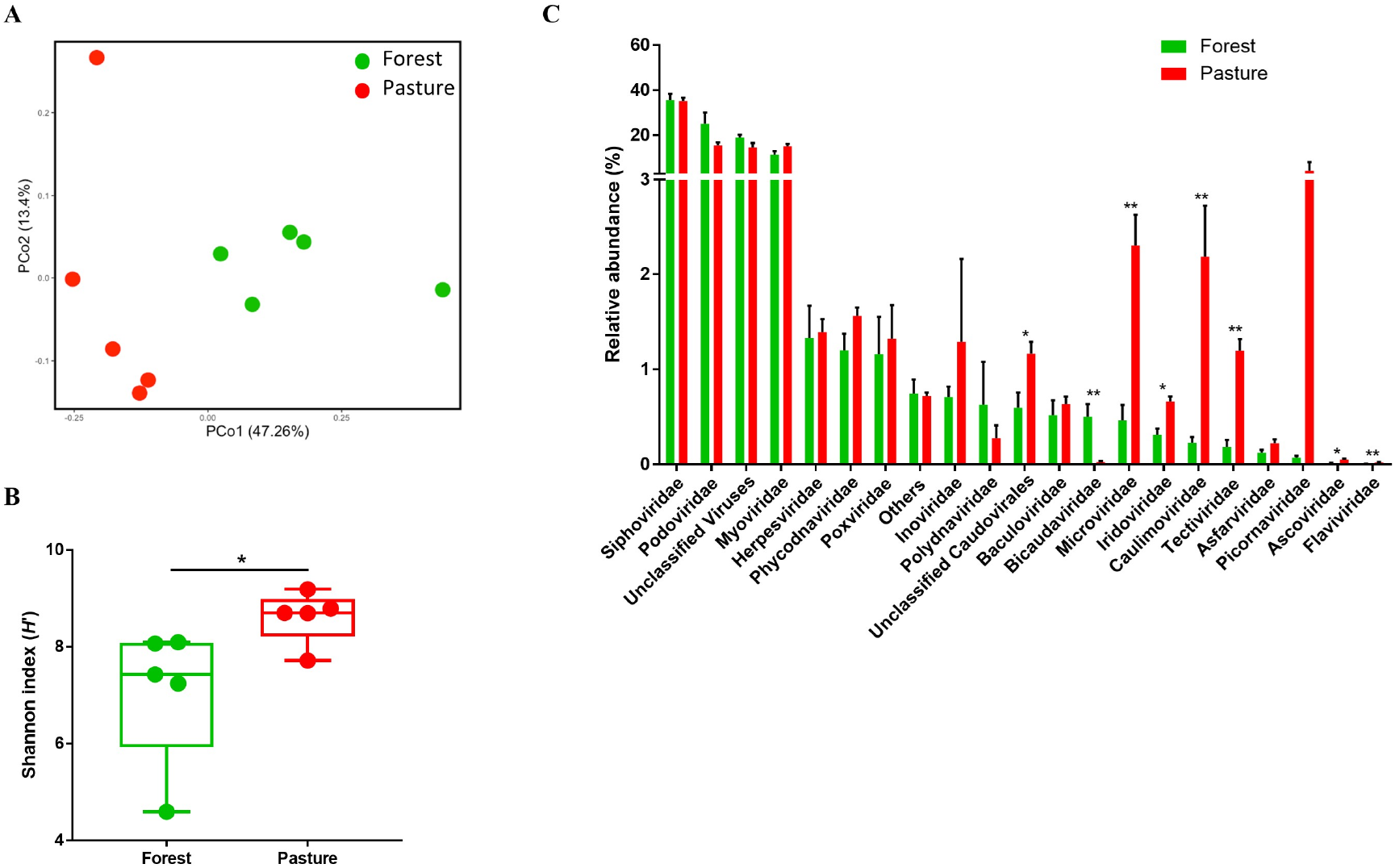
Virome community structure in the metagenomes obtained from the Amazon forest (green) and pasture (red) soils. (A) Ordination plot from principal coordinate analysis showed that forest viral composition is distinct from that of pasture. (B) Alpha diversity as estimated by Shannon index is significantly higher in pasture compared to forest. (C) Relative abundances of viral communities are estimated at family level in forest and pasture metagenomes. Error bars represent standard error of the mean (S.E.M). Symbols * and ** indicate significance values of *P* < 0.05 and *P* < 0.01, respectively, which were calculated using Mann-Whitney test with 1,000 permutations.

## DISCUSSION

By applying a multifactorial strategy of deep sequencing (mean 636 million reads per sample) and a machine learning approach, we compared taxonomic differences and generated metabolic maps with differential representation of genes involved in the cycling of nutrients and greenhouse gases in two land use types in the Amazon basin: a pristine forest and a 38 years old pasture. Importantly, we reported diversity and compositional differences of viral communities in forest and pasture, which remained unstudied for these tropical ecosystems. The alteration of bacterial and archaeal compositions and diversities following deforestation, which we observed in this study, mirrors previous observations (5, 14–16). These microbial communities are compositionally in strong concordance with protein-coding (KO) genes, suggesting that soil samples share many of these species, with associated KO genes and metabolic processes. In addition, our study demonstrated that ecosystem conversion results in increased homogenization of protein coding genes (KO), which may be explained by the reduced substrate heterogeneity in the pasture dominated with only two grass species. These shifts in biochemical pathways imply changes in the microbial physiological strategies. Importantly, these differences in metagenomes between forest and pasture are strongly associated with variation of energy and carbohydrate metabolisms; two processes of importance to carbon and nitrogen cycling. While most of the major functional characteristics are maintained in both ecosystems, relative abundances of many specific genes in these two processes substantially differed, indicating the potential for differences in physiological status and nutrient cycling between forest and pasture.

Plants are major contributors of organic carbon to soils through rhizodeposition, and microbial communities utilize these carbon compounds (40). The metagenomes of pasture and forest show signatures of resulting resource differences. The pasture soil is rich in cellulosic litter from grasses, and one of the two grass species that dominates in pasture is *Urochloa brizantha*, which stores nutrients including starch materials in their rhizomes (41). Along with animal dung in pastureland, these carbon compounds potentially represent rich sources of fermentable carbohydrates. In the pasture, the increased occurrence of genes involved in starch and sucrose metabolism, and the transport of mono- and di-saccharides suggest that microbes may rely more on exogenous saccharides than other substrates. This may explain why pasture metagenomes have elevated counts of genes for glycolysis, TCA cycle and acetate fermentation. In comparison, the forest metagenome was enriched in genes involved in gluconeogenesis. Higher abundance of pentose phosphate pathway (PPP) genes in forest metagenomes may provide flexibility to glycolysis/gluconeogenesis by funneling glyceraldehyde-3-P, an intermediate of the PPP, into these pathways. Here, we report that forest metagenomes had higher biosynthetic capacity of essential amino acids of oxaloacetate, pyruvate and PPP families, which potentially favors the growth of the bacterivorous and detritivorous animal component of the soil ecosystem. On the other hand, higher gene abundances for amino acid biosynthesis in the α-ketoglutarate family were demonstrated in microbial communities of pastureland. An amino acid of this family, glutamate, is the precursor for chlorophyll synthesis (42), which may support the increased capacity for photosynthesis and carbon fixation of microbial communities in pastureland. This is in agreement with previous findings of increased microbial biomass for the same pasture evaluated in our study (43).

In contrast to the pasture, the forest soil is a more oligotrophic environment, and along with a lower concentration of substrates, there is also a higher diversity of C compounds due to the complex secondary chemistry of a myriad of forest vegetation (44, 45). These complex compounds are degraded by soil microorganisms into a few central aromatic intermediates (*e.g*., benzoate, catechol, etc.) through peripheral catabolic pathways that are subsequently channeled into the β-ketoadipate pathway by *ortho* ring cleavage (46). Among others, nitrate-respiring bacteria, dominated by pseudomonads, cleave the benzene-ring of these aromatic compounds (47–50), which leads to the generation of TCA cycle intermediates. Here, it is noteworthy that *Pseudomonas* spp. and genes related to the degradation of these compounds are mostly enriched in forest soils, suggesting the maintenance of TCA cycle by replacing oxaloacetate and pyruvate that are removed for the biosynthesis of glucose and amino acids. The overrepresentation of genes associated with glyoxylate metabolism in forest may also contribute to carbon anaplerosis by supplying malate and its subsequent conversion to oxaloacetate. Together, the carbohydrate metabolic processes in forest microbial communities not only would maintain a better trade-off between glycolysis and gluconeogenesis but also provide metabolic plasticity for energy harvest and anaplerosis, a strategy that may be critical for the oligotrophic forest soil environment.

Prominent differences imposed by ecosystem conversion in the Amazon involve pathways related to energy metabolism and observed differences involve metabolisms of methane and nitrogen, which support previous observations (6, 15, 16, 25, 27, 51). Our data demonstrate that *Euryarchaeota*, where known methanogens belong (25), and several known KO genes associated with methanogenesis are elevated following deforestation. Increased capacity of glycolysis and acetate fermentation may fuel the methanogenesis in pasture, which subsequently may increase the efficiency of fermentation by scavenging acetate and producing carbon dioxide. In contrast, methane-consuming microbial taxa and associated genes involved in this pathway have lower abundances in the pasture, particularly due to the loss of proteobacterial methanotrophs including *Methylomonas*, *Methylobacterium* and *Methylocella*. In addition to differences in methane cycling, our results highlighted differences in pathways of nitrogen cycling. Higher gene content to nitrify nitroalkanes in the forest suggests that nitroalkanes may serve as sole sources of nitrogen, carbon, and energy in its microbial communities (52, 53). Our results also suggest that nitrifying archaeal phylum *Thaumarchaeota* and several species of it are more frequently distributed in forest soils, which was also shown in our previous study using functional marker genes (24). In addition, we observed higher abundance of several species of *Alphaproteobacteria* in forest soils known to be associated with steps of the nitrogen cycle, such as *Nitrobacter* sp. LIP in nitrification, and *Rhodoplanes* sp. HMD01017 in denitrification. Higher capacity for energy-yielding denitrification in forest soils and other oligotrophic habitats leads to the increased production of nitrous oxide and nitrogen gases (Tiedje, 1988), which is evident in the Amazon (55).

Soil physicochemical properties not only impact on microbial composition, but also the rate of biogeochemical processes (57, 58). Previous reports suggest that temperature is a key factor that regulates many terrestrial processes including soil respiration (59), nitrification (60), denitrification (61) and methane emission (62). In addition, the C/N ratio has substantial impacts on nitrogen mineralization and nitrification rates (63–65), and therefore on denitrification. Deforestation by slash-and-burn processes causes a variety of changes in chemical factors (i.e., nutrients) that depend on land use type and depth, time since conversion, soil treatment and amendments, and subsequent land use (66). To the extent that available nitrogen declines, there is likely selective pressure on soil microbes to scavenge atmospheric nitrogen. This probably explains the higher capacity of pasture for nitrogen fixation, consistent with previous works showing increased abundance of the *nifH* gene (22, 67). The higher C/N ratio and the lability of carbon sources, evidenced by overrepresentation of genes associated with starch and sucrose metabolism in the pasture, are suggestive of reduced capacity for nitrification and therefore denitrification. On the other hand, a meta-analysis by Rustad et al. (2001) reported that warming in the range 0.3–6.0°C increased soil respiration rates by 20% and net N mineralization rates by 46%. Temperature induced increased respiration would lead to a depletion of oxygen in the terrestrial ecosystem, which subsequently would trigger anaerobic energy-yielding processes such as fermentation, denitrification and methanogenesis. This observation has important implications. The temperature difference between our study sites (24.86°C in forest versus 27.14°C in pasture) (15) is similar to what is predicted by 2050 in the Amazon (68). While both soils are sources of greenhouse gases (nitrous oxide, nitric oxide, CO_2_, CH_4_), increasing temperature have higher potential for the generation of these greenhouse gases from both ecosystems. Therefore, our study in two adjacent ecosystems along temperature gradient provides an opportunity to explore the long-term impact of climate-driven increase in temperature on ecosystem processes in the Amazon soils.

Finally, our deep sequencing approach allowed us to obtain a glimpse of the Amazon virome. Viruses are reservoirs of many protein-coding genes of prokaryotic hosts (69), with the potential to influence nutrient cycling, and the adaptation and evolution. For instance, *Pseudomonas* phage PAJU2, a lysogenic phage (70), was estimated to be the most abundant virus in forest soil and its relative abundance is about 8-fold higher compared to the pasture soils, suggesting its important role in maintaining the population size and activity of ecologically important *Pseudomonas* hosts in forest soils. Importantly, the enormous microbial and viral deposits in the forest soil may have novel infectious strains or virulent strains of known environmental microorganisms. Clearing the pristine forest may pose a potential threat to human and animal health. Following the arrival of Europeans in the New World, millions of Native Americans in or near the Amazon basin were killed due to outbreaks of deadly infections, notably smallpox. Our results showed that *Molluscum contagiosum* virus and *Vaccinia* virus, two members of *Poxviridae* family are overrepresented in forest soils. Such soils could preserve cowpox, and even the otherwise extinct smallpox virus. Increased incidences of poxvirus infections in cattle and humans have been reported near the Amazon Region of western Colombia (71) and throughout Brazil (72), coinciding with the increased rate of deforestation in the Amazon. *Wolbachia* phage WO is also overrepresented in forest soils. As one of most widespread reproductive parasites in the biosphere (73), *Wolbachia* influences the reproductive capacity of hosts, including mosquitos, and lytic WO phage is associated with the reduction of *Wolbachia* densities and reproductive parasitism (74). For this and other reasons (e.g., increased puddle formation due to tree stump removal and cattle trampling), forest-to-pasture conversion may drive mosquito-borne disease events, including the recent zika outbreak, and continuous malaria threats in Brazil and Peru near the deforested areas (75). Increased malaria also followed deforestation in Malaysian Borneo (76). Knowledge of the disease ecology of deforestation is still rudimentary, underscoring the need for increased surveillance of potential disease outbreaks near deforested regions.

### Caveats

We acknowledge caveats for our study. First, the trade-off between deep sequencing and low sample size prevented a complete exploration of spatial variability in metagenome profiles. However, previous sequencing and empirical studies covering broader spatial ranges in the pristine Amazon and deforested areas are in line with the current observations, suggesting that higher sample sizes will further consolidate the observed differences between forest and pasture. Second, while a vast majority of the sequence reads passed the quality filtering, only 13.4% of the sequence reads were annotated, a percentage similar to other soil shotgun metagenomic studies (77, 78). The biology of a vast majority of microbial species remains unknown, limiting full annotation potential genes in metagenomes regardless of database used. Third, metagenomes analyzed in the study provide a snapshot of the functional characteristics of microbial communities at a given time, which may not illuminate important phenomena such as disturbance, recovery, and succession. Fourth, there is a lack of well-curated databases and well-established methods for annotating terrestrial virome at species level. In general, viral sequences are poorly represented at species level in public databases. There are some platforms that are designed and optimized for the detection of viruses of specific kinds such as VirSorter (79) and vConTACT (80) for bacterial and archaeal viruses, and ViromeScan (81) for eukaryotic and human viruses. While there are methods that are recently developed for assembled (VirMap) (82) and unassembled (FastViromeExplorer) (83) metagenomic sequences, none of them has been validated for soil virome. Therefore, in order to be consistent with the algorithm and database used for annotation of prokaryotic taxonomy, we also used RefSeq database to annotate viral taxonomy through the MG-RAST method that provides species level resolution.

## Materials and Methods

### Site description and sampling

The study was performed at the Amazon Rainforest Microbial Observatory (ARMO), located at Fazenda Nova Vida in the State of Rondonia, Brazil (10°10′18.71″S, 62°47′15.67″W) (5). This location has one of the highest rates of deforestation in the Brazilian Amazon (84), representing the last four decades of man-made ecosystem alterations occurring in this region. Detailed information about the vegetation and soil properties of our study sites have been described elsewhere (6, 9). The primary forest is a typical wet “Terra Firme” forest with the majority of the trees yet to be identified, while the pasture was established in 1972 after a slash-and-burn procedure followed by aerial seeding of the fast-growing grasses *Urochloa brizantha* and *Panicum maximum*. The sampling strategy and sample processing were described in detail elsewhere (5, 25). Briefly, in each of the forest and pasture ecosystems, five nested quadrats were previously established within a 1-hectare plot. These two sampling plots are 3.66 km apart. For this study, sampling points were located at the corners of each quadrat along a transect, and soil cores were collected with a 10 cm-depth by 5 cm-diameter corer after the removal of the litter layer. The five soil cores for each land use type were collected in 2010, frozen on the spot, transported on dry ice to the laboratory and stored at −80°C until DNA extractions. Soil physicochemical properties were analyzed at the Laboratorio de Fertilidade do Solo, Department of Soil Sciences, University of Sao Paulo. At the time of sampling, *in situ* measurements of soil pH averaged 3.96 and 4.24, while soil temperature averaged 24.8°C and 27.2°C for forest and pasture, respectively.

### Soil DNA extraction and sequencing

Soil DNA extraction was performed with the PowerLyzer Powersoil DNA isolation kit (Mobio Inc., Carlsbad, CA). Five DNA extractions were performed per soil sample and pooled for metagenomic library preparation with the TrueSeq DNA kit (Illumina Inc., San Diego, CA). Sequencing was performed on the Illumina HiSeq platform to generate 6.4 billion of paired-end reads of 150bp.

### Sequence processing and annotation of metagenomes

The Amazon soil metagenomic dataset is publicly available and deposited in the Rapid Annotation using Subsystems Technology for Metagenomes (MG-RAST) database (85). A link for a total of 83 MG-RAST IDs of five forest and five pasture samples is provided in Meyer et al. (2017). Quality filtering of the raw sequences was performed using the default criteria as set by the MG-RAST pipeline (i.e. trimming of low-quality bases, removal of artificial replicate sequences and filtering of sequences with greater than 5 ambiguous bases). As part of the MG-RAST pipeline version 3.2, paired-end reads were joined using the fastq-join algorithm. Single end reads with 150 bp length that could not be joined were retained. Quality filtered sequence reads were searched against the Kyoto Encyclopedia of Genes and Genomes (KEGG) (86), and RefSeq (87) databases for functional and taxonomic (bacterial, archaea and viruses) annotations, respectively. The annotations were conducted at an e-value cutoff of 1e-5, a minimum identity cutoff of 60%, and a minimum alignment length cutoff of 15 (default parameters) for the annotations of metagenome sequences. Following the annotations, the datasets were exported in biom file format and merged by QIIME 1.8.0 (88) for the downstream analyses.

### Data analysis

Rarefaction plots were generated to check whether sampling effort was sufficient. We then compared the relative abundances of major bacterial, archaeal and viral taxa, and functional categories of genes (KEGG level 2) between forest and pasture ecosystems. Alpha diversities were calculated using species (or gene) richness and the Shannon index (*H’*). Bray-Curtis dissimilarity was used to calculate the pairwise dissimilarities and perform principal coordinate analysis (PCoA) between metagenomes. To test the hypothesis that forest and pasture have differential metagenome profiles, PCoA plots were generated using the species level abundance dataset of bacteria and archaea, where functional categories at KEGG level 2 were fitted as vectors. Here we used two *R* packages-‘*ape*’ (89) and ‘*vegan*’ (90) to calculate the association of each ecosystem with different functional attributes of the bacterial and archaeal communities.

### Statistical Analysis

To determine whether differences in the relative abundances of major taxonomic and functional categories between ecosystems were statistically significant, the Mann-Whitney test with 1000 permutations was used. Performing the same statistical test, we compared the alpha diversities, and pairwise taxonomic distances between samples. Analyses of similarities (ANOSIM) tests were performed to assess whether metagenome compositions were significantly different across the terrestrial ecosystems.

We employed the DESeq2 method (91), a negative binomial Wald test, to identify microbial and viral species, and protein-coding genes at KEGG level 4 (KEGG orthology, KO) that were significantly affected by forest-to-pasture conversion. The negative binomial distribution is particularly useful for metagenomic datasets, where observations have high variation among biological replications, and sample variance is typically greater than the sample mean (91). This method also calculates fold-difference in gene abundance, which is represented as the log_2_-fold-change (log_2_FC). A score of zero indicates that the KO gene or species has statistically the same proportional abundance in metagenomes of both land use types. A positive value for a given gene or species indicates overrepresentation in forest metagenomes compared to pasture, while a negative value indicates the reverse pattern. We also employed a supervised classifier, Random Forest using the *R* package ‘*randomForest*’ (92), to classify a subset of soil species and protein-coding genes that are discriminatory between land uses. To estimate the accuracy of the model, a subset of the samples was randomly partitioned as a training set to build the model, which was then used to classify the remaining samples as test set (cross-validation). With leave-one-out cross validation, suitable for small sample sizes, the average estimated generalization error of the classifier set was calculated and compared with the baseline error for random guessing. We calculated the ratio of baseline error and estimated error, and considered this ratio of two or higher for a good classification, meaning the Random Forest classifier performs at least two-fold better than random guessing. Additionally, this classifier calculates an importance score for each feature (i.e., species or protein-coding gene) by estimating the increase in estimated error when a feature is ignored. In our analysis, we considered 0.0001 as the cutoff for importance score. The use of Random Forest in conjunction with DESeq2 was conservative, in that more genes and species passed only the DESeq2 screen, and fewer passed both. Together, both of the statistical tests increased the robustness in identifying the differentially abundant species and genes between ecosystems with their levels of importance in the ecosystems.

QIIME 1.9.0 and GraphPad Prism 7.00 were used for performing statistical analyses and visualizing the results.

## Availability of data and materials

Metagenomic sequences for all samples used in this study are available at the Joint Genome Institute IMG/M database under the following accession numbers: 1080879 to 1080888.

## ACKNOWLEDGMENTS

The authors thank the owners and staff of Agropecuaria Nova Vida for logistical support and permission to work on their property. We are grateful to Siu M. Tsai, Rebecca Mueller, Fabiana Paula, Babur Mirza, and Wagner Piccinini for assistance with fieldwork.

## SUPPLEMENTAL MATERIAL

**FIG. SF1. Rarefaction curves of bacterial (A), archaeal (B), viral (C) taxonomic and protein-coding (KO) (D) gene distributions of metagenomes obtained from Amazon forest and pasture soil samples.** Species richness was used as a function of sequencing depth for forest and pasture samples. Green and red lines indicate forest and pasture samples, respectively.

**FIG. SF2. Relative abundances of bacterial (A) and archaeal (B) communities at phylum level in the metagenomes of Amazon forest (green) and pasture (red).** Taxa that are shown in each of the bacterial and archaeal domains represent over 98% of their taxonomic sequences. The most abundant phylum *Proteobacteria* is broken down into five classes. Error bars represent standard error of the mean (S.E.M). *P*-value is calculated using Mann-Whitney test with 1,000 permutations and symbols * and *** indicate significance values of *P* < 0.05 and *P* < 0.001, respectively.

**FIG. SF3. Relative abundances of functional categories at KEGG level 2 in the metagenomes obtained from the Amazon forest (green) and pasture (red).**Error bars represent standard error of the mean (S.E.M). Symbols * and *** indicate significance values of *P* < 0.05 and *P* < 0.001, which were calculated using Mann-Whitney test with 1,000 permutations.

**FIG. SF4. KEGG pathway map of differentially abundant genes observed in forest and pasture metagenomes for amino acid metabolism of (A) oxaloacetate acid, (B) pentose phosphate pathway, (C) pyruvate, and (D) alpha-ketoglutarate.** Only genes that passed both selection criteria for DESeq2 (BH-adjusted *P* < 0.05) and Random Forest (importance score > 0.0001) methods are represented. Arrows indicate the enzyme-mediated steps of the pathway, with KEGG EC numeric classification representing the reaction being catalyzed. A green arrow means that the abundance of a gene encoding KEGG EC enzyme was higher in the forest, while red means higher in the pasture. Black arrows indicate genes with similar abundances between forest and pasture metagenomes.

**FIG. SF5. KEGG pathway map for xenobiotic metabolism involved in benzoate, dioxin, and polycyclic aromatic hydrocarbon degradation.** Only genes that passed both selection criteria for DESeq2 (BH-adjusted *P* < 0.05) and Random Forest (importance score > 0.0001) methods are represented. Arrows indicate the enzyme-mediated steps of the pathway, with KEGG EC numeric classification representing the reaction being catalyzed. A green arrow means that the abundance of a gene encoding KEGG EC enzyme was higher in the forest, while red means higher in the pasture. Black arrows indicate genes with similar abundances between forest and pasture metagenomes.

**FIG. SF6. Relationships of protein-coding (KO) gene compositions with soil physicochemical factors.** Vector fitted principal coordinate analysis of protein-coding genes with vectors representing different soil factors. Each vector points to the direction of an increase in the gradient for the corresponding variable and length proportional to the correlation between ordination and variable. The circles (green, forest; red, pasture) represent the relative positions of protein-coding genes observed in a sample. The significances (*P*-values) of the vectors were calculated based on 999 random permutations of the data. Only soil physicochemical factors that were estimated to be significantly (*P* < 0.01) associated with PCo1 are shown. Bray-Curtis metric was used for estimating distances between different samples.

**FIG. SF7. Relationships of viral composition with bacterial (A), archaeal (B), and protein-coding gene (C) compositions across forest and pasture metagenomes.** The relationships of compositional similarities are visualized using the principal coordinate loadings for the first axes (PCo1). The circles (green, forest; red, pasture) represent the relative positions of each soil taxonomic community. Bray-Curtis metric was used for estimating distances between different samples.

**TABLE ST1. A complete list of bacterial (sheet A) and archaeal (sheet B) lineages at species level that discriminate the metagenomes between forest and pasture soils.** DESeq2 (BH-adjusted *P* < 0.05) and Random Forest (importance score > 0.0001) methods were employed to calculate the discriminatory prokaryotic and viral species between forest and pasture metagenomes. Random Forest is a supervised classifier that classifies a subset of protein-coding genes that are discriminatory between ecosystems. This classifier calculates an importance score for each feature (i.e., protein-coding gene) by estimating the increase in generalized error when feature is ignored. In our analysis, we considered 0.0001 as the cutoff for importance score. On the other hand, DESeq2 employs negative binomial distribution to identify the differentially abundant protein-coding genes, where log_2_FC value is calculated to demonstrate the degree of variations in abundances for each protein-coding gene. Here, the positive and negative log_2_FC values indicate the genes that are differentially abundant in forest and pasture metagenomes, respectively. The lfcSE stands for standard error of log_2_FC estimate. *P*-value is calculated by the Wald statistic, which is the ratio of log_2_FC and its standard error. *P*-values were adjusted for multiple comparisons using Benjamini-Hochberg False Discovery Rate, and adjusted *P* < 0.05 was used for statistical significance.

**TABLE ST2. A complete list of protein-coding genes at KEGG level 4 that discriminate the metagenomes between forest and pasture soils.** DESeq2 and Random Forest methods were employed to calculate the protein-coding genes that differentiate between forest and pasture metagenomes.

**TABLE ST3. A complete list of viral lineages at species level that discriminate the metagenomes between forest and pasture soils.** DESeq2 (BH-adjusted *P* < 0.05) and Random Forest (importance score > 0.0001) methods were employed to calculate the discriminatory prokaryotic and viral species between forest and pasture metagenomes.

